# Front propagation and arrival times in networks with application to neurodegenerative diseases

**DOI:** 10.1101/2022.01.04.474911

**Authors:** Prama Putra, Hadrien Oliveri, Travis Thompson, Alain Goriely

## Abstract

Many physical, epidemiological, or physiological dynamical processes on networks support front-like propagation, where an initial localized perturbation grows and systematically invades all nodes in the network. A key question is then to extract estimates for the dynamics. In particular, if a single node is seeded at a small concentration, when will other nodes reach the same initial concentration? Here, motivated by the study of toxic protein propagation in neurodegenerative diseases, we present and compare three different estimates for the arrival time in order of increasing analytical complexity: the linear arrival time, obtained by linearizing the underlying system; the Lambert time, obtained by considering the interaction of two nodes; and the nonlinear arrival time, obtained by asymptotic techniques. We use the classic Fisher-Kolmogorov-Petrovsky-Piskunov equation as a paradigm for the dynamics and show that each method provides different insight and time estimates. Further, we show that the nonlinear asymptotic method also gives an approximate solution valid in the entire domain and the correct ordering of arrival regions over large regions of parameters and initial conditions.

## 1 Introduction

We live in a connected world where people, goods, information, and diseases travel from one region to the next. In the years of COVID-19, a particularly dramatic example of this propagation phenomenon is, of course, the transmission of the coronavirus from a single seed location (Wuhan, China), to the rest of the world through an intricate network of local and global travel routes [1]. A similar phenomenon also appears in a much smaller system, the human brain, where some toxic proteins associated with neurodegenerative diseases like Alzheimer’s or Parkinson’s are believed to originate in a single region [2] and are transported to the rest of the brain through the so-called structural connectome, the network of axonal pathways connecting different regions of the brain [3, 4].

From a mathematical perspective, both phenomena can be understood as the propagation of an autocatalytic process on a network, and the main question is to understand its overall dynamics [5, 6]. For instance, if a process such as a disease starts at a seed location, how long will it take to appear at other locations, and then develop through a full-scale invasion, leading to a global pandemic for a disease, or to dementia for the brain? The *arrival-time problem* consists in determining the time it takes for a quantity of interest to reach a certain level at each node. It has been shown that the scaling of arrival times greatly depends on the coupling dynamics between nodes [7]. Here, we restrict our attention to the important case of linear coupling between nodes, typically given by the graph Laplacian, which is used both for the progression of diseases in global networks [8, 9] and in the modeling of neurodegenerative diseases [10].

### 1.1 Previous work

There has been a great deal of attention to the problem of arrival times, especially within the context of epidemiology. There are essentially two different approaches from a modeling perspective. The process is either seen as a stochastic random walk problem on a graph with some form of autocatalytic expansion at a node [11], or as the discretization of a continuum reaction-diffusion process on the network. While many authors view the stochastic process as the ground truth and the continuum perspective as a mean-field approximation [12, 13, 14], we take the view that the continuum system is a model of the dynamical process, independently of any underlying discrete process [15, 16]. The central question is then to obtain meaningful estimates about its dynamics rather than justifying the use of a mean-field approach. Therefore, we place ourselves at the continuum deterministic level and do not address further similar models based on stochastic processes.

Within the framework of continuous processes, there are three different approaches to obtain estimates for the arrival time in the case of a nonlinear process on a network. First, one can simplify the topology of the problem and consider the equivalent front propagation on a homogeneous tree or a complete graph. Then, by using a discrete version of the continuous front problem that arises in nonlinear parabolic equations, one can obtain a velocity of propagation [17, 18]. While the mathematical theory for these systems is appealing and has led to many interesting results in their own rights [19], it only applies to network with very particular structures. However, for the systems we are interested in, the network topology is particularly important and has a strong influence on the arrival times. Hence, we will not consider such methods here.

A second approach consists in studying the linearized system around an equilibrium solution. Then the concentration at each node can be obtained as the solution of this linear system for a given initial condition. From this solution, the arrival times are given as the solution of a transcendental equation at each node [20]. This approach is particularly good for small initial concentrations and large diffusion as the linearization provides a decent estimate of the solution. Based on that approach, a suitable ansatz for the nonlinear behavior can be introduced to improve the arrival time estimates [21]. However, the fact that the arrival time is given by a numerical solution of a transcendental equation does not provide much insight into the dynamics. Moreover, without a good prior guess, this solution may be difficult to obtain.

The third approach consists in looking at the propagation between two nodes while neglecting all other connections and, again, linearize this system. We can then extract an estimate of the time it takes for the second node to reach the same level as the initial condition in the seed node [12]. This arrival time provides a notion of distance between nodes [22] and the arrival time between two distant nodes can then be approximated as the shortest path between these nodes according to this notion of *effective distance* [22]. An improvement based on the contribution due to multiple paths can also be obtained at the cost of extra computations [13]. This notion of distance on a network is particularly powerful as it gives the network an extra structure from which many estimates can be obtained.

At this point, it is important to reflect on why we need an arrival-time estimate in the first place. If the exercise is to obtain a numerical value, then a direct simulation of the system of nonlinear ordinary differential equations is easy to implement and provides an easily computable solution to the problem, even for very large systems. Typically, the solutions obtained by solving transcendental equations in the linear cases or by using the effective distance over multiple paths are not as precise and cannot be obtained as easily, both in terms of implementation and in terms of computational cost. Hence, the goal here is not to obtain a method to compute the arrival times, as this problem can be solved numerically, but to obtain meaningful estimates and approximations. There are two main reasons why approximations and estimates may be valuable. First, we want to answer basic questions in terms of the parameters: how long does it take for the first seed to reach neighboring nodes? How does the first phase of the process depend on the parameters or topology of the system? Once the initial invasion takes place, can we estimate the velocity of invasion? How long does it take for the system to be fully invaded? Second, we may need a decent analytical approximation of the solutions themselves so that it can be used to understand other coupled processes. We will show that both goals can be achieved by using a combination of approximate distances and asymptotics.

### 1.2 Fisher-KPP on networks

The simplest model for an invasion process on an undirected connected network is a discrete version of the celebrated Fisher-Kolmogorov-Petrovsky-Piskunov (Fisher-KPP) reaction-diffusion equation [23]. Consider a connected and undirected network *G* = (*V*, *ℰ*) with a collection of *N* nodes *V* representing regions of interest and edges *ℰ* representing connections between these regions. If *p*_*i*_ = *p*_*i*_(*t*) is the quantity of interest in a region *i*, modeled as a node in a network, and evolving with time *t*, then on this network, the dynamics is governed by a system of *N* ordinary differential equations

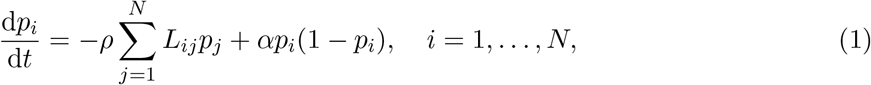

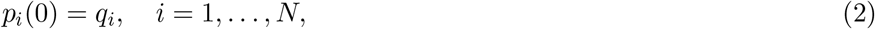

where *ρ, α* are positive and respectively depict the diffusion constant and growth constant, and *q*_*i*_ ∈ [0, 1] for all *i*. The matrix **L** is the symmetric graph Laplacian [24] obtained from the symmetric weighted-adjacency matrix **A** as

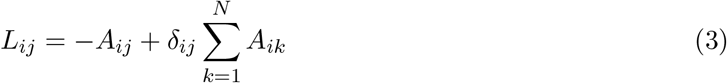

where *δ*_*ij*_ is the Kronecker delta (*δ*_*ij*_ = 1 if *i* = *j*, 0 otherwise). The graph Laplacian models diffusion processes on a network and encodes the connection between regions: *L*_*ij*_ = 0 if there is no connection between different nodes *i* and *j*. For comparison between different networks, we further assume that the graph Laplacian has been rescaled so that 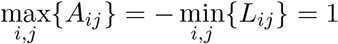.

Here, we are using the so-called *standard graph Laplacian* in contrast to one the many normalized versions of the Laplacian which are often preferred in the mathematical study of graphs [25]. However, as discussed in [26], our choice of Laplacian is the only one that, in the absence of a reaction term, satisfies the two fundamental properties required to model diffusion: overall mass conservation, and the Fick’s property that no transport takes place on the network if the concentrations at each node are equal.

For the *canonical arrival-time problem*, the system is seeded at a single node *s* so that *q*_*s*_ = *β* and *q*_*i*_ = 0, *i* ≠ *s*. Then, a simple question is to obtain the value of the *arrival times τ*_*i*_ = *τ*_*i*_(*μ*) such that *p*_*i*_(*τ*_*i*_) = *μ* (with *β* ≤ *μ <* 1 so that *τ*_*i*_ ≥ 0 and finite with the equality *τ*_*i*_ = 0 attained only if *i* = *s* and *μ* = *β*). Unless specifically stated, we will mostly study the case where *μ* = *β* and the problem is to know the times at which other nodes reach the same initial value as the seeding node. The *generalized arrival-time problem* consists of a system with an arbitrary initial condition. Then, let 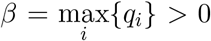 and the corresponding question is to obtain the value of the arrival times *τ*_*i*_ = *τ*_*i*_(*μ*) such that *p*_*i*_(*τ*_*i*_) = *μ* (with *β* ≤ *μ <* 1 so that *τ*_*i*_ ≥ 0 for all *i*).

Since the network is undirected and connected, the system only supports two equilibrium states when *α* and *ρ* are non-vanishing: the *healthy state* defined by *p*_*i*_ = 0 for all *i*, and the *toxic state* defined by *p*_*i*_ = 1 for all *i*. The healthy state is unstable while the toxic state is stable. The dynamics in the absence of transport (*ρ* = 0) follows locally a logistic equation, with a sigmoid solution. Hence, for all values 0 *< β <* 1 of the seed, the system will evolve towards the toxic state, which implies that the arrival times *τ*_*i*_ are all positive and finite.

By construction, the graph Laplacian has one zero eigenvalue. The dynamics is therefore mostly controlled by *λ*_2_, the smallest non-zero eigenvalue of **L**. Indeed, we can define two main regimes of interest for this system as shown in Fig. 1: the *diffusion-dominated regime*, defined by *ρλ*_2_*/α* ≫1, and the *growth-dominated regime*, for which *ρλ*_2_*/α* ≪ 1.

**Figure 1:**
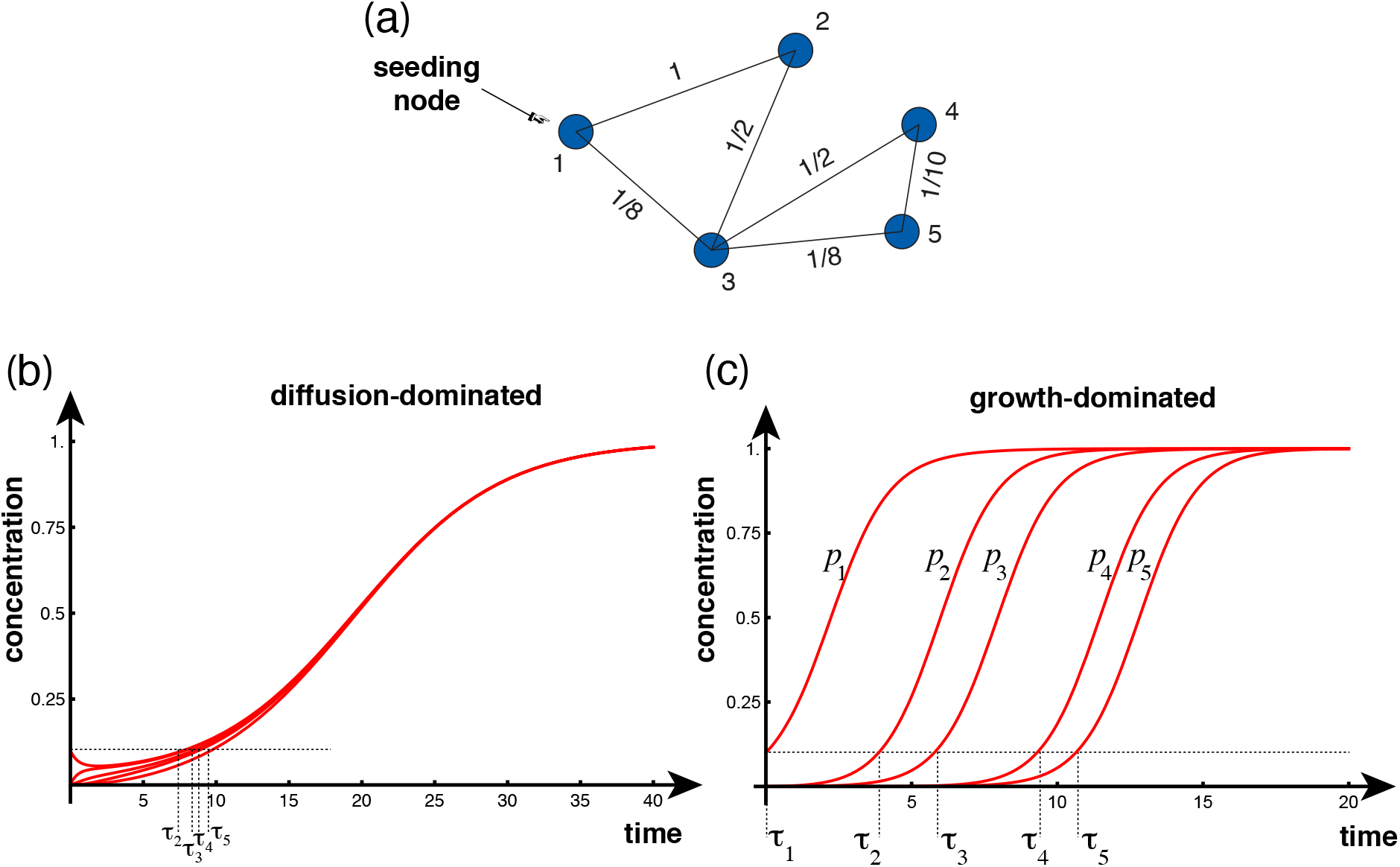
Example of dynamics on a 5-node network (a). Initially, only the first node is seeded (*p*_1_(0) = 1*/*10). In the diffusion-dominated case (b: *α* = 1*/*5, *ρ* = 1), all concentrations quickly approach 1*/*10 before feeling the effect of the growing exponential in unison. For the growth-dominated case, after growth at the first node, the second node is invaded and then all other modes are subsequently invaded in a front like progression (c: *α* = 1, *ρ* = 1*/*100).

As a guiding example, consider the 5-node network shown in Fig. 1 with weighted adjacency matrix

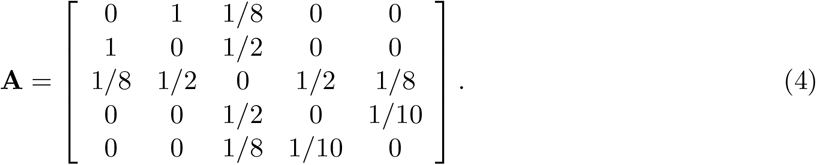

The graph Laplacian for this network, obtained from (3), has smallest non-zero eigenvalue *λ*_2_ ≈ 0.228.

Intuitively, one expects that in the diffusion-dominated regime, shown in Fig. 1b, the concentration will rapidly become uniform over the network, as predicted by a purely diffusive mechanism. Then, a local slow exponential growth will take place more or less simultaneously at all nodes.

The growth-dominated regime is somewhat more interesting from a dynamical point of view. In this case, the concentration at the seeded node will increase while diffusion takes place. The effect of the diffusion term is then to seed other connected regions in which the concentration will then also increase as shown in Fig. 1c. In this case, the system is systematically invaded, node per node, and the question is to understand the dynamics of this cascade by using the arrival times as road markers.

## 2 A linear analysis

### 2.1 General method

The first naive approach to the arrival-time problem is to linearize the system around the healthy state and integrate it. In this case, the system reduces to

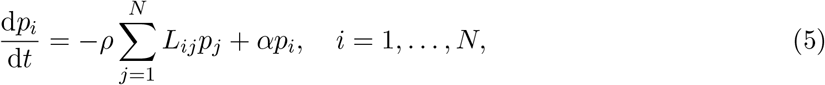

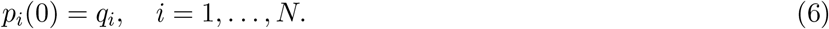

A compact solution to this problem is obtained in terms of matrix exponentials. Let **p**(*t*) = (*p*_1_, …, *p*_*N*_), **q** = **p**(0) = (*q*_1_, …, *q*_*N*_), and **M** = −*ρ***L** + *α***1**, where **1** denotes the identity matrix. Then, the solution is given by

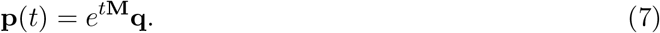

Since **L** is symmetric, there exists a complete set of orthonormal eigenvectors **v**^(*i*)^ ∈ ℝ^*N*^ associated with the ordered (but not necessarily distinct) eigenvalues *λ*_*i*_, *i* ∈ {1, …, *N*} so that

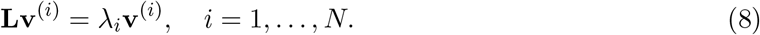

Further, by construction, we have *λ*_1_ = 0 with **v**^(1)^ = (1, …, 1) and, since the network is connected, 0 *< λ*_*i*_ ≤ *λ*_*j*_ for all 1 *< i < j* [25]. Then, the linear solution can be written explicitly as

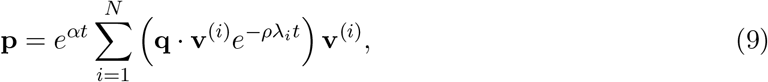

and the *general linear arrival time* 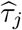 is given by the solution of the equation

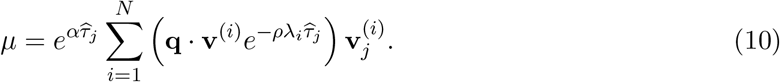

If we restrict our attention to the *canonical linear arrival time*, we can further simplify the system by using *q*_*r*_ = *βδ*_*rs*_ to obtain

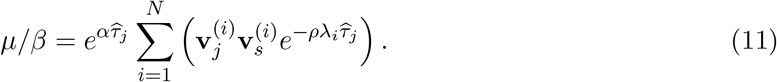

A comparison of the linear solutions with the numerical solutions for our 5-node network is given in Fig. 2bc. A few general features of the method emerge. First, the approximation given by the linear arrival times improves as *β* = *μ* decreases. Indeed, as a consequence of the stable manifold theorem, there exists an open set in phase space **p** ∈ ℝ^*N*^ such that the linear solution is a faithful approximation of the solution. Therefore, as *μ*→ 0 with *μ >* 0, this approximation converges to the exact value as the nonlinear terms becomes negligible. This behavior is illustrated Fig. 2 where we compute the error made as a function of the seed size:

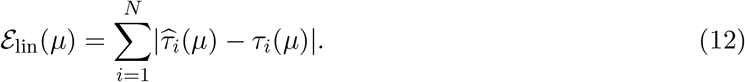

**Figure 2:**
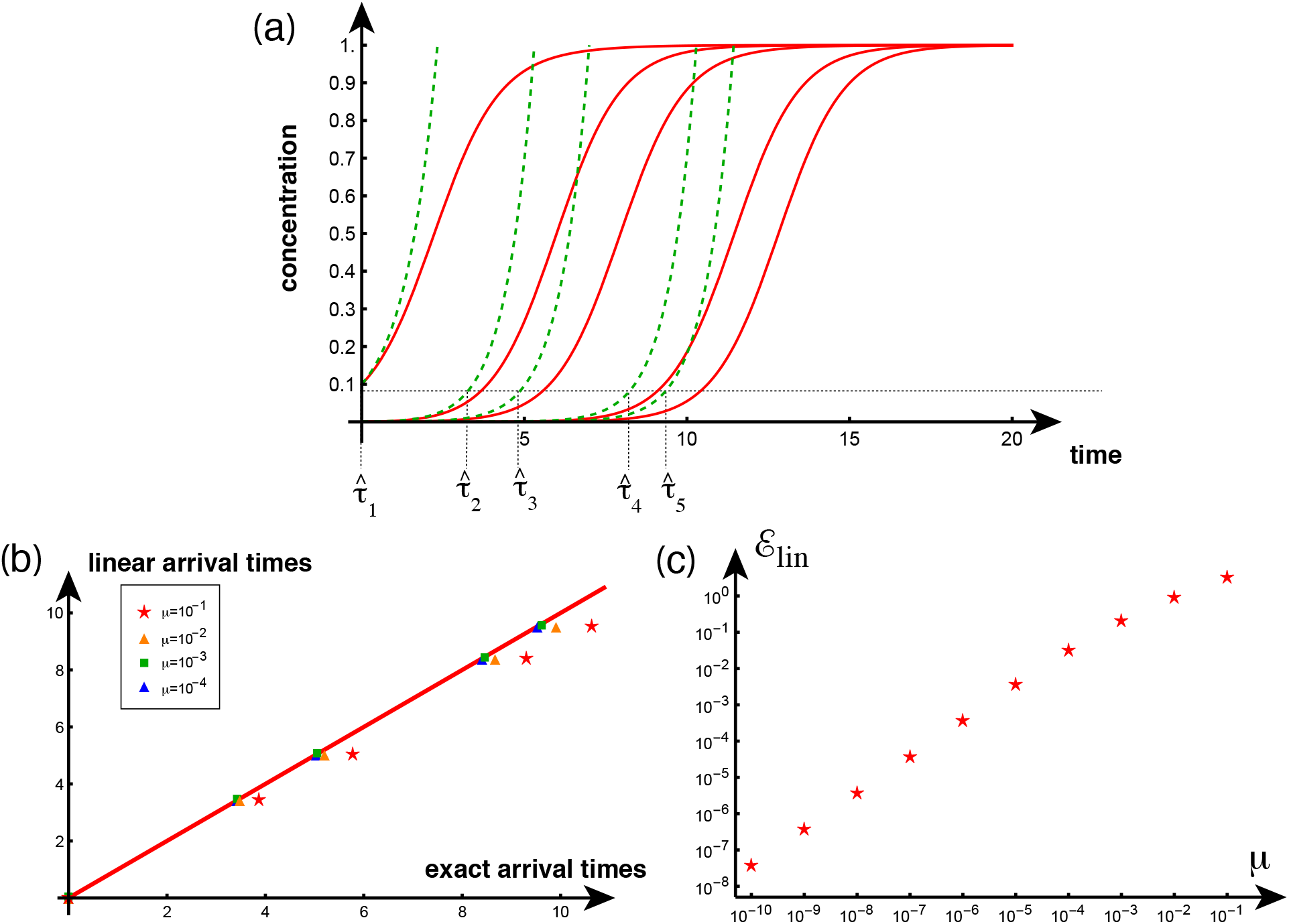
Comparison between the exact (red) and linear solutions (dashed, green) for the 5-node network (a: *α* = 1, *ρ* = 1*/*100, *β* = 1*/*10). Exact arrival times vs linear arrival times for various initial seeding (and *β* = *μ*) (b) and error between exact and linear arrival times (c) as a function of initial seeding shows good convergence, as expected.

Second, since the nonlinear terms of (1) act to reduce the value of the concentration in time, and (5) neglects these terms, the solutions of the linear equations are always strictly larger than the actual concentrations (apart from the zero arrival time at the seeding node). Hence, the linear estimates provide a strict lower bound for the arrival times.

Third, to compute the linear arrival times we need to solve *N* individual transcendental equation of the form (10), one for each arrival time. Solving these equations can prove to be numerically challenging without a decent first guess, a problem that we address in the next section.

Fourth, this method is often overlooked at the favor of more involved graph-based measures based on the superposition of multiple paths. However, the exponential matrix solution (7) naturally combines the sum over all paths, due to the properties of the graph Laplacian, as expected from a diffusion operator. Therefore, the linear arrival times constitutes a robust universal method valid for small seed concentrations to obtain arrival times as further demonstrated in the next sections.

### 2.2 Diffusion-dominated propagation

The linear analysis is sufficient to understand fully the diffusion-dominated case. Indeed, if diffusion dominates, then the early dynamics is governed by the slowest dynamics in (11), which is the term involving exp(−*ρλ*_2_*t*). Therefore on a typical time scale *t* ∼ 1*/*(*ρλ*_2_), the concentration at each node tends to the value *μ/N*. On larger time scales, the effect of the growing exponential is felt and each node behaves, in unison, as a single node subject to the dynamics

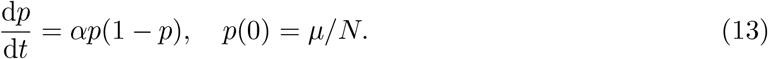

The solution of this equation is

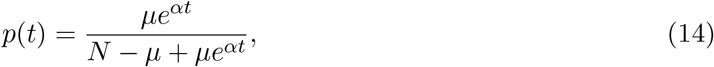

with an arrival time at all nodes given by

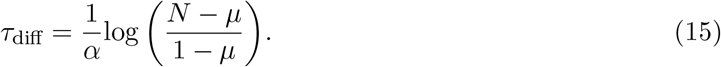

We see in Fig. 3 that both estimates give an excellent approximation for the overall behavior of the system and the arrival time.

**Figure 3:**
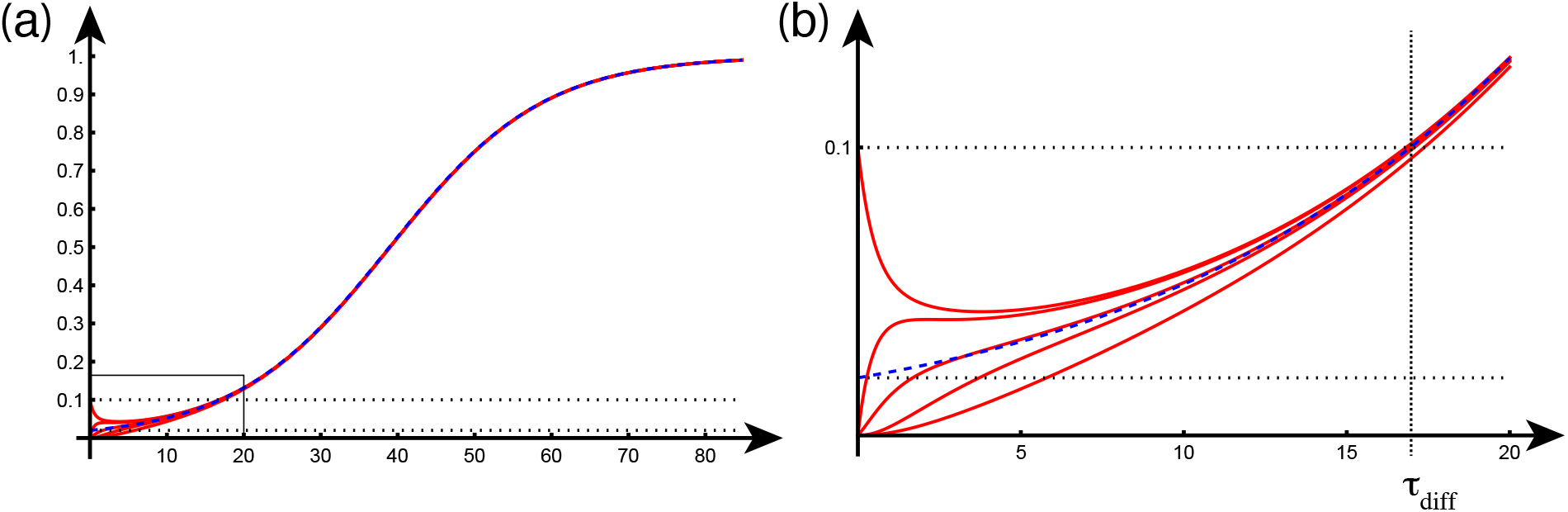
Diffusion-dominated case for the 5-node network with *α* = 1*/*10, *ρ* = 1, *p*_1_(0) = *μ* = 1*/*10. We see that the dynamics of a single node with initial condition *p*(0) = *μ/N* (dashed curve) provides an excellent approximation of the entire solution (a) and, looking at the close-up in the grey rectangle, for the arrival time as well (b).

There are two interesting aspects to this dynamics. First, in the diffusion-dominated case, after an early period, the concentration increases at each node simultaneously. Hence there is no clear distinction between the different curves, as small changes in the initial conditions can change the order at which each node reaches the critical threshold. Hence, we do not expect to see a well-established ordering of the nodes by arrival times. Second, we see that the concentration at the seeding node decreases initially through diffusion. From the general equations, the slope of the concentration at the seeding node and at the initial time *t* = 0 is given by

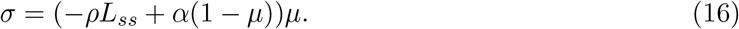

Therefore, the concentration at the seeding node will initially decrease if *ρL*_*ss*_ *> α*(1 − *μ*), which will occur if the diffusion constant is large enough. The critical threshold *ρL*_*ss*_ = *α*(1 − *μ*) provides another key criterion for the initial dynamics.

For the rest of the paper, we will focus on the richer growth-dominated case, for which we assume, without loss of generality, that *α* is of order one and *ρ* is a small parameter.

## 3 The Lambert distance

An second approach is to define a notion of distance between nodes by considering the time it takes for an initial seed to propagate to a neighboring node while ignoring all other nodes in the network. Consider two nodes *i* and *j* connected by an edge with weight *A*_*ij*_ ≠ 0. Neglecting all other nodes in the network, we consider the linear approximation (5) with surrogate graph Laplacian, corresponding to the two-node subnetwork of node *i* and *j*, given by

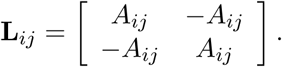

Using the eigenvectors of **L**_*ij*_ and (11) with *μ* = *β* yields an equation for the arrival time at node *j*, from an initial seeding at node *i* (or vice-versa), given by

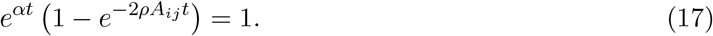

Expanding for small *ρ*, we obtain

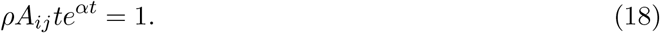

The solution of this equation can be expressed in terms of the Lambert function *W*_0_ (defined so that the real solution of *te*^*t*^ = *z* is *t* = *W*_0_(*z*)):

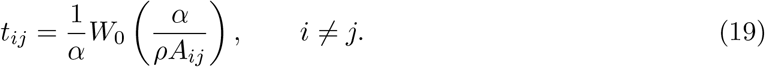

We note that since the network is undirected, we have *A*_*ij*_ = *A*_*ji*_ which implies that *t*_*ij*_ = *t*_*ji*_.

We refer to *t*_*ij*_ as the *Lambert edge distance*. From this pairwise distance between neighboring nodes, we define the *Lambert distance W*_*ij*_ as the shortest path with respect to the Lambert edge distance between two nodes *i* and *j*. Explicitly, let Γ_*ij*_ = (*γ*_0_, *γ*_1_, …, *γ*_*n*_) with *γ*_0_ = *i* and *γ*_*n*_ = *j* be this shortest path, then

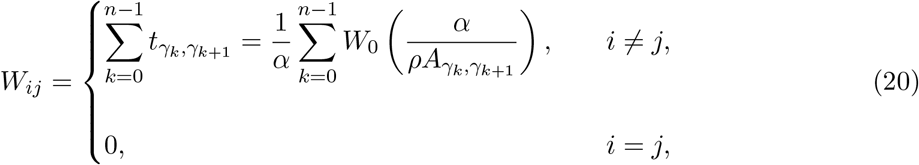

which defines the *Lambert distance matrix* **W**. By construction, this distance is a metric on the network (as *W*_*ii*_ = 0 ∀*i, W*_*ij*_ = *W*_*ji*_ ∀*i, j* and *W*_*ij*_ ≤ *W*_*ik*_ + *W*_*kj*_ ∀*i, j, k*).

The Lambert distance provides a useful estimate for the arrival times. To take into account the fact that, in general, the critical concentration may be different from the initial concentration, *μ* ≠ *β*, we define the *self-time t*_*ii*_ to be the time at which a local initial concentration *β*, at node *i*, reaches *μ* in the absence of any connection. That is, the time *t*_*ii*_ at which the solution to

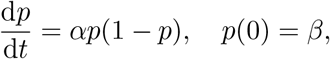

satisfies *p*(*t*_*ii*_) = *μ*. The solution of this problem is

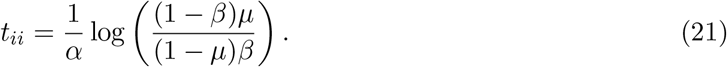

In the canonical case of a single seed at node *i*, the *Lambert arrival times*, at every node *j*, are defined by

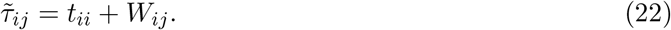

Since both *W*_*ij*_ and *t*_*ij*_ are symmetric, it follows that 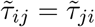 It is useful to note that the term *W*_*ij*_, in (22), is computed by (20), which implicitly assumes that *μ* = *β* via (17)-(19), and that the self-time, *t*_*ii*_ of (21), vanishes when *μ* = *β* a priori. The Lambert arrival times (22) can then be understood as the sum of the time it takes for node *j* to reach the same concentration as the seed node *i* and the time for node *j* to reach concentration *μ* starting with concentration *β*. Equivalently, it can also be interpreted as the sum of the time it takes for node *i* to reach concentration *μ* and the time it takes for the concentration *μ* at node *i* to reach node *j*.

This estimate misses three contributions, which prevents it to be an upper or lower bound. First, any node before the target node is connected to other nodes (including previous nodes). Hence, the quantity of interest is diluted away from the target node which reduces the arrival time. Second, the target node may also receive contributions from other paths, resulting conversely in overestimated arrival time. Third, the estimate is based on a linearization of the equations, hence it misses saturating effects that increase the arrival time. Hence, unlike the linear arrival times, the Lambert arrival times (22) do not provide a general bound, neither upper or lower, for to the true arrival times.

Yet, as shown next, the Lambert method provides a remarkably good estimate despite its simplicity. It has a few notable features. First, the entries of the Lambert distance matrix (20) are independent of the initial concentration as the initial concentration acts only to perturb the Lambert arrival time (22) via the self-arrival (21). The Lambert distance, between two distinct nodes, is therefore an intrinsic property of the system. It implies that the initial seed concentration does not appreciably impact the overall propagation. Second, the Lambert method highlights the role of the diffusion in the arrival-time problem and, in particular, shows that diffusion has only a weak influence on this quantity. For instance, for *ρ/*(*αW*_*ij*_) ≈ 0.01, an increase of diffusion by a factor 10 only changes the arrival times by a factor 2. Third, the Lambert arrival times can be used as a a natural first guess to find the linear times and greatly improve the convergence of root-finding methods applied to (11).

From the Lambert arrival times, we can obtain simple expressions for the front solution. This is done by assuming that the dynamics after the arrival time is controlled by the local dynamics so that, after this time, the effect of the graph Laplacian is neglected. More specifically, assume that node *i* is chosen as the seeding location, with initial seed *p*_*i*_(0) = *μ*, then the dynamics at node *j* is given by

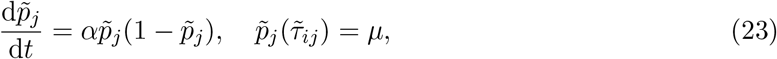

where 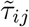 is the Lambert arrival time (22). The *Lambert solution*

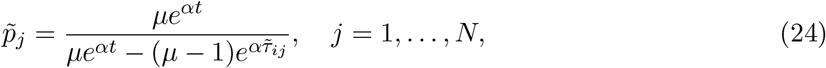

is therefore an approximation of (1).

The Lambert edge distance (19), Lambert distance (20), and Lambert solution (24) can be readily illustrated using the exemplar 5-node network of Fig. 1. The matrices corresponding to the Lambert edge distance and Lambert distance, respectively, are:

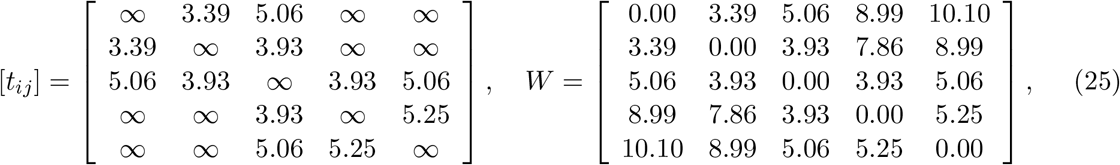

where we have used the symbol ∞ to characterize pairs of nodes that are not connected by en edge (and hence have an infinite Lambert edge distance). an example of the corresponding Lambert solutions and the arrival times is given in Fig. 4. The Lambert solutions, and Lambert arrival times, are seen to be an improvement over both the linear solutions, and arrival times, and for capturing the dynamics over the entire domain.

**Figure 4:**
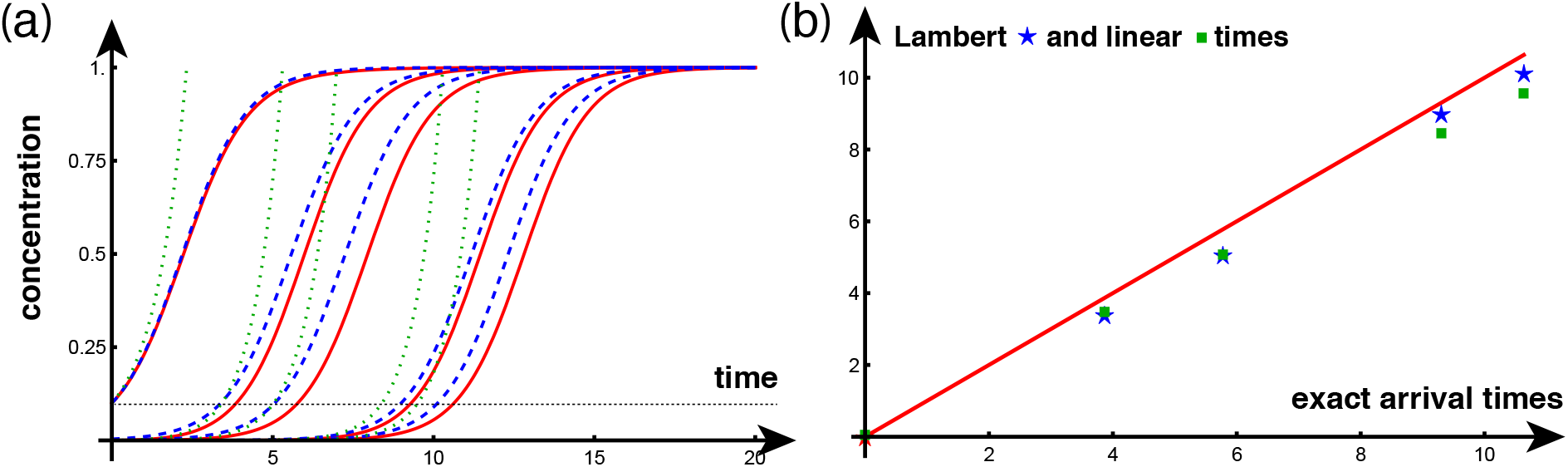
a. Exact (red), Lambert (blue, dashed), and linear (green, dotted) solutions with *α* = 1, *ρ* = 1*/*100, *p*_1_(0) = *μ* = *β* = 1*/*10. b. Lambert arrival times (blue stars) compared to linear arrival times (green squares).

Finally, we introduce the notion of *step distance*. The step distance is a vector, ***γ***, whose *j*^th^ entry is the number of edges along the shortest path from the node of initial seeding to the node *j*. For the 5-node network example, of Fig. 1, with seeding at node 1 we have ***γ*** = (0, 1, 1, 2, 2). Thus, for example, this particular ***γ*** indicates that shortest path, with respect to the Lambert metric established by **W**, between seed node 1 and target node 3 is comprised of 2 edges.

### 3.1 Multiple seeding locations

The Lambert distance assumes that a single node *i* has been seeded and gives the arrival times *W*_*ij*_ at node *j*. In the case where multiple seeding locations are given at time 0, the method can be easily adapted. Indeed, assume that a general initial condition **p**(0) = **q** is given, with 0 ≤ max_*i*_{*q*_*i*_} *< μ* ≪ 1. The current task is to derive a closed-form expression for an approximate arrival time 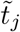, in a target node *j*, such that 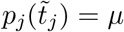. Fix a target node *j* and consider a seed node, *i*, such that 0 *< q*_*i*_. Following the previous analysis, we use (22), with *β* = *q*_*i*_, to obtain

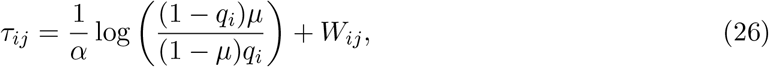

which estimates the arrival time at node *j* due to the seed at node *i*. Now, we construct an approximation of the contribution to the concentration at node *j* due to the seeding at node *i* by

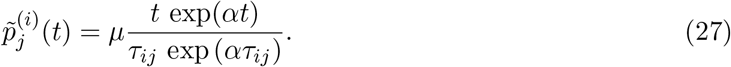

Summing over the partial contributions from the multiple seeding sites yields an approximation of the total concentration at node *j* given by

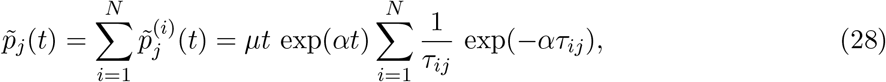

which reaches the level *μ* at time

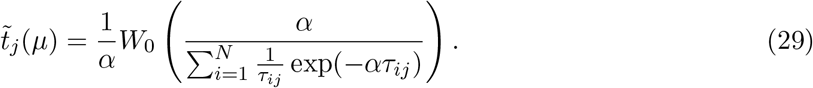

Just as in the case of a single seeding site, the estimate (29) can be used as an initial guess to compute the linear arrival times for the case of multiple seeding locations.

### 3.2 Discussion and comparison with other notions of distance

The idea to use the Lambert function as an approximation for the arrival times was first proposed in the seminal paper [12] in conjunction with an effective edge distance replacing (19) by

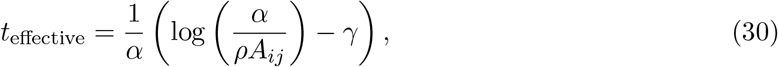

where *γ* is the Euler constant. Surprisingly, despite showing better fit to the data (See Fig. 2 in [12]), the Lambert distance was disregarded by the authors and the focus of the community moved to the notion of the effective distance based on the logarithmic edge time (30). Following the influential work of Brockman, Helbing and co-authors, who introduced the fundamental notion of a metric based on logarithmic edge times [13, 22], the effective distance gained further popularity, probably due to a familiarity with the logarithmic function. Despite some attempts to relate both notions [21], there is no meaningful limit in which the two distances coincide as can be observed directly from the graphs of these two function shown in Fig. 5. The difference between the two estimates creates a systematic bias and we found that the effective distance does not perform as well as the Lambert distance, especially on long paths.

**Figure 5:**
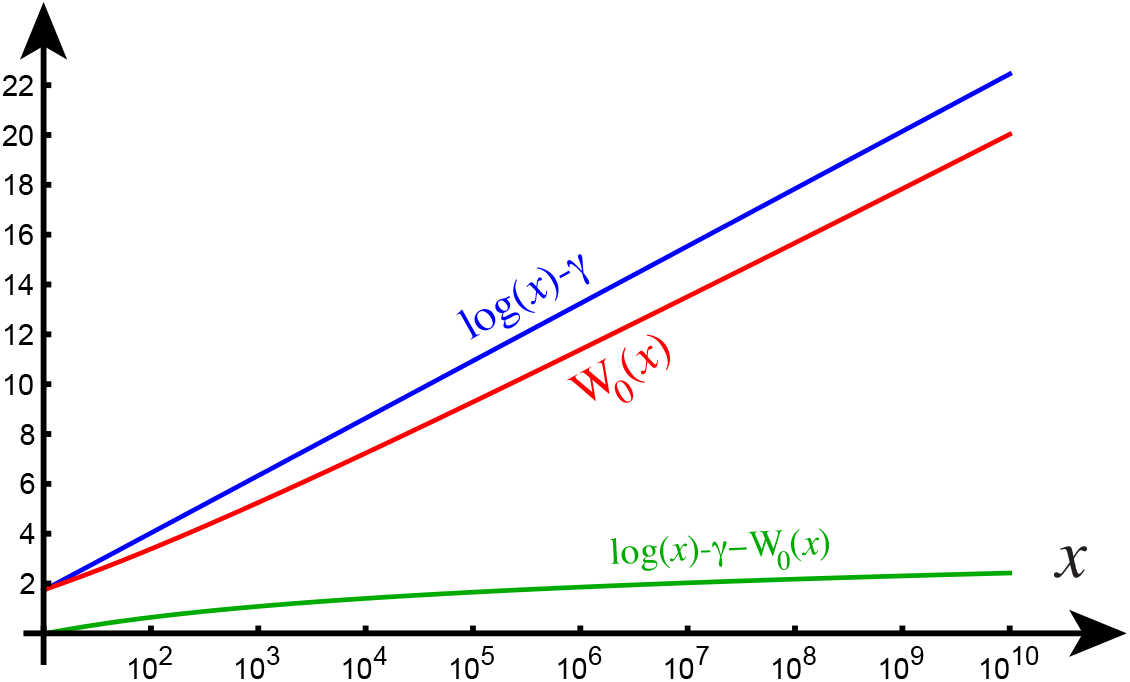
Comparison of the effective edge time (blue) and the Lambert edge time (red) as a function of their argument in the growth-dominated case. We observe that the use of the logarithm function for the distance creates a systematic overestimate of the arrival time of order one in the case of two nodes. In a network, this difference (shown in green) adds to every single edge along a path and causes a systematic error.

We also note that the presence of the Euler constant in (30) is related to the stochastic approach of [12] who showed that, within the probabilistic framework, arrival times follow a Gumbel distribution. Within the context of a deterministic model, however, there is no a priori justification for this constant despite the fact that it lingers in the literature like a vestigial appendage.

## 4 Nonlinear approximations

The estimates obtained so far are useful but they do not fully exploit all the information that can be obtained from the nonlinear system. In particular, these method do not take into account the coupling between the nonlinear sigmoid response encapsulated in the nonlinear term and the diffusion from nearby nodes given by the graph Laplacian. In the growth-dominated case, we have a natural small parameter that can be used for asymptotic computation.

Assume, by a simple rescaling in time and without loss of generality, that *α* is of order one and that *ε* = *ρ* ≪ 1 is a small parameter. A naive asymptotic method that systematically expands all solutions in terms of *ε* and solve the resulting equations order by order fails to provide any useful results as it only provides a small correction of the exponential behavior which is unbounded in time.

To circumvent this difficulty, we present an alternative asymptotic method that captures important features of the solutions and improves the computation of the arrival times. We consider the canonical case and start with the solution with *ε* = 0 given by

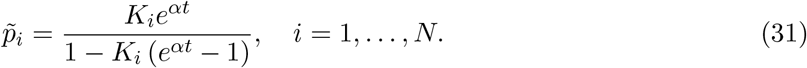

At this point, we resist the urge to set the arbitrary constant from the initial conditions. Rather, we use a nonlinear version of the method of variation of constants^1^

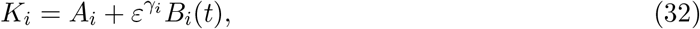

where *γ*_*i*_ is the step distance between seed node and node *i* as defined in Section 3). With this ansatz, we can solve iteratively for the unknown function *B*_*i*_ for the subsequent subsets of nodes with increasing distance from the seeding node up to the furthest node given by the maximum of ***γ***.

To lowest order *O* (*ε*^0^), the solution at the seeding node with initial condition *p*_*s*_(0) = *β* is simply the original unperturbed solution

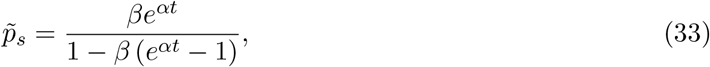

and *A*_*i*_ = 0 for all *i* ≠ *s*. To order *O* (*ε*^1^), representing all nodes *i* at a distance one from the seeding node, the differential equation for *B*_*i*_ is

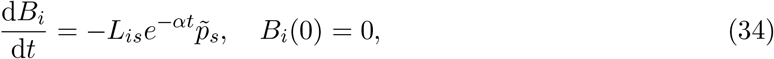

that can be easily integrated to obtain

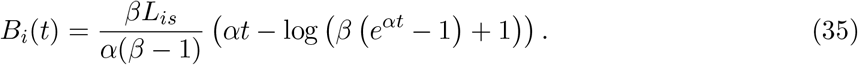

The solutions for nodes at distance two from the seeding node can then be obtained iteratively with nodes at distance *d* only depending of the solutions given by nodes at distance *d* − 1. More formally, if we define the matrix

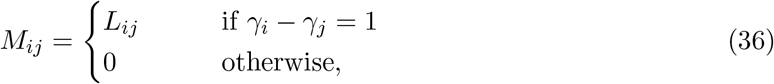

and the variables

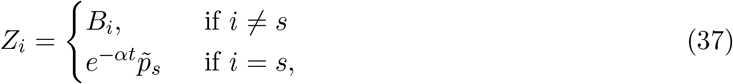

then, these new variables are solutions of the system

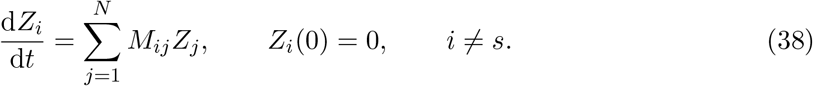

The asymptotic solutions, for the 5-node network of Fig. 1, are shown in Fig. 6 together with the computations of the arrival times.

**Figure 6:**
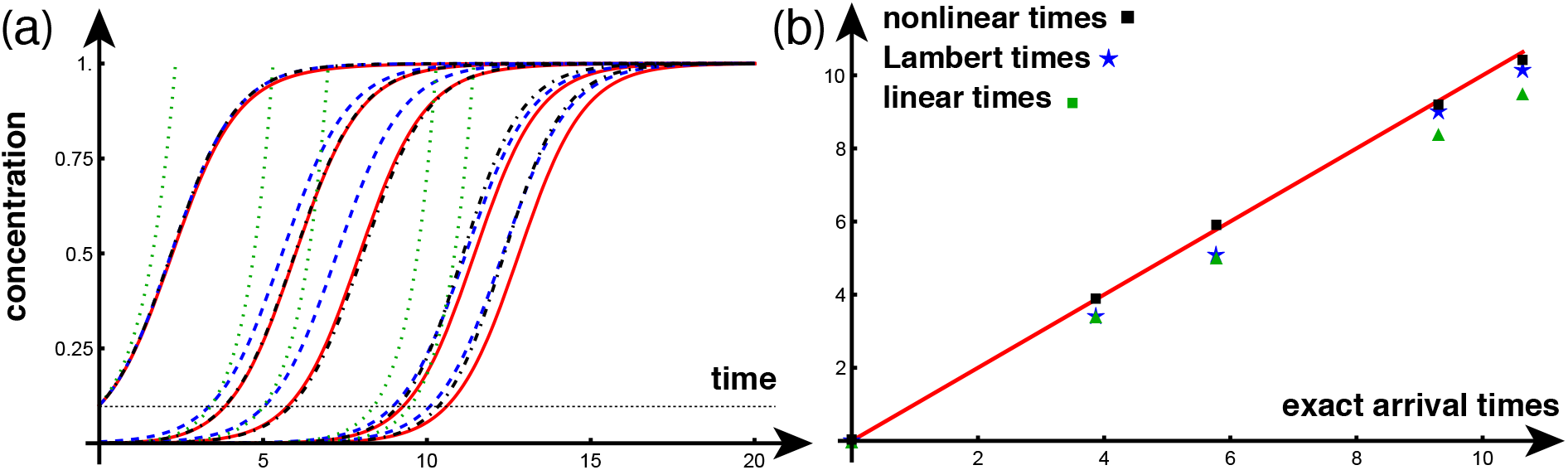
a. Exact (red), Lambert (blue, dashed), linear (green, dotted), and asymptotic (black, dash-dotted) solutions with *α* = 1, *ρ* = 1*/*100, *p*_1_(0) = *μ* = 1*/*10. b. Asymptotic arrival times (black squares), versus Lambert arrival times (blue stars) versus linear arrival times (green squares).

Remarkably, the solution of (38) can be obtained explicitly in terms of the polylogarithm function Li_*d*_ where *d* is the distance to the seeding node (for node at distance 1, the polylogarithm reduces to the natural log as found in (35)). Once substituted in (31), these *asymptotic solutions* satisfy both the initial conditions and the asymptotic behavior for long time. They naturally take into account the contributions from all paths of length *d* rather than just a single path. Hence, they improve the Lambert solution, albeit at the cost of increasingly long expressions better carried through symbolic manipulations in the privacy of one’s office. They do not include the contributions due to paths of different lengths or the effect of dilution away from a node. Both of these effects could be, in principle, included in the matrix **M** by adding the corresponding couplings and introducing higher-order corrections, but at the cost of an increase in complexity that may not be warranted.

## 5 Application to neurodegenerative diseases

In this section, we apply the arrival time methods to the study of propagation of misfolded toxic protein species, transported on a structural connectome. Indeed, it is now well appreciated that neurodegenerative diseases such as Parkinson’s and Alzheimer’s are associated with toxic proteins forming large aggregates. Starting in an initial region, these toxic proteins can systematically invade the brain through the axonal pathways known as the structural connectome.

A *structural connectome* and its weighted adjacency matrix **A**, can be obtained by tractographic reconstruction of axonal bundles from diffusion tensor magnetic resonance images. The structural network of a patient brain is then defined by a set of vertices reflecting cortical and subcortical regions of interest and a set of edges representing the axonal bundles that connect these regions. The structural networks used here are obtained from 426 patient data graphs part of the Budapest Reference Connectome v3.0 [29, 30] and generated using a multi-resolution Lausanne anatomical parcellation [31, 32]. The connectome networks are freely available [31] at five levels of resolution and we will use averaged connectome with the lowest (*N* = 83) and highest (*N* = 1015) resolution [33]. These networks have edges with weight *A*_*kj*_ between nodes *k* and *j*, defined as the ratio of the number of axonal fibers between the nodes and the mean fiber length squared. With these brain networks in hand, different mathematical models of misfolded protein propagation have been systematically used and validated to study key aspects of neurodegenerative diseases [34, 35, 33]. For instance, Alzheimer’s disease is associated with the formation of large aggregates of amyloid-beta and tau proteins. The accumulation of toxic tau proteins is particularly important to understand the disease as it is associated with regions of atrophy and cognitive dysfunction. Its progression has two key features of relevance for the present work: the toxic load originates in localized areas of the brain around the entorhinal cortex [23] and a systematic pattern of progression has been identified through post-mortem histopathology, suggesting that the progression is in the growth-dominated regime with a small diffusion constant [36, 37].

Therefore, we model the propagation of toxic tau protein through the network model of Fisher-KPP and used our methods to understand the dynamics.

### 5.1 Time scales of disease progression

In Fig. 7, we show the solution of the Fisher-KPP equation on the small scale connectome with 83 nodes for particular initial conditions and parameter values. We are now in a position to reflect on the different methods and extract from our analysis important features that answer the question we started with: How do local spreading and global propagation depend on the system’s parameters? In the growth-dominated case, we see a clear separation of time scales. The *local time scale:*

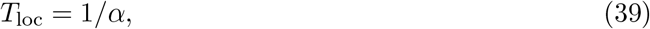

governs the local increase of any seed. Indeed, an initial concentration *β <* 1*/*2 reaches locally a concentration of 1/2 at a time

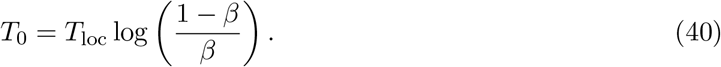

**Figure 7:**
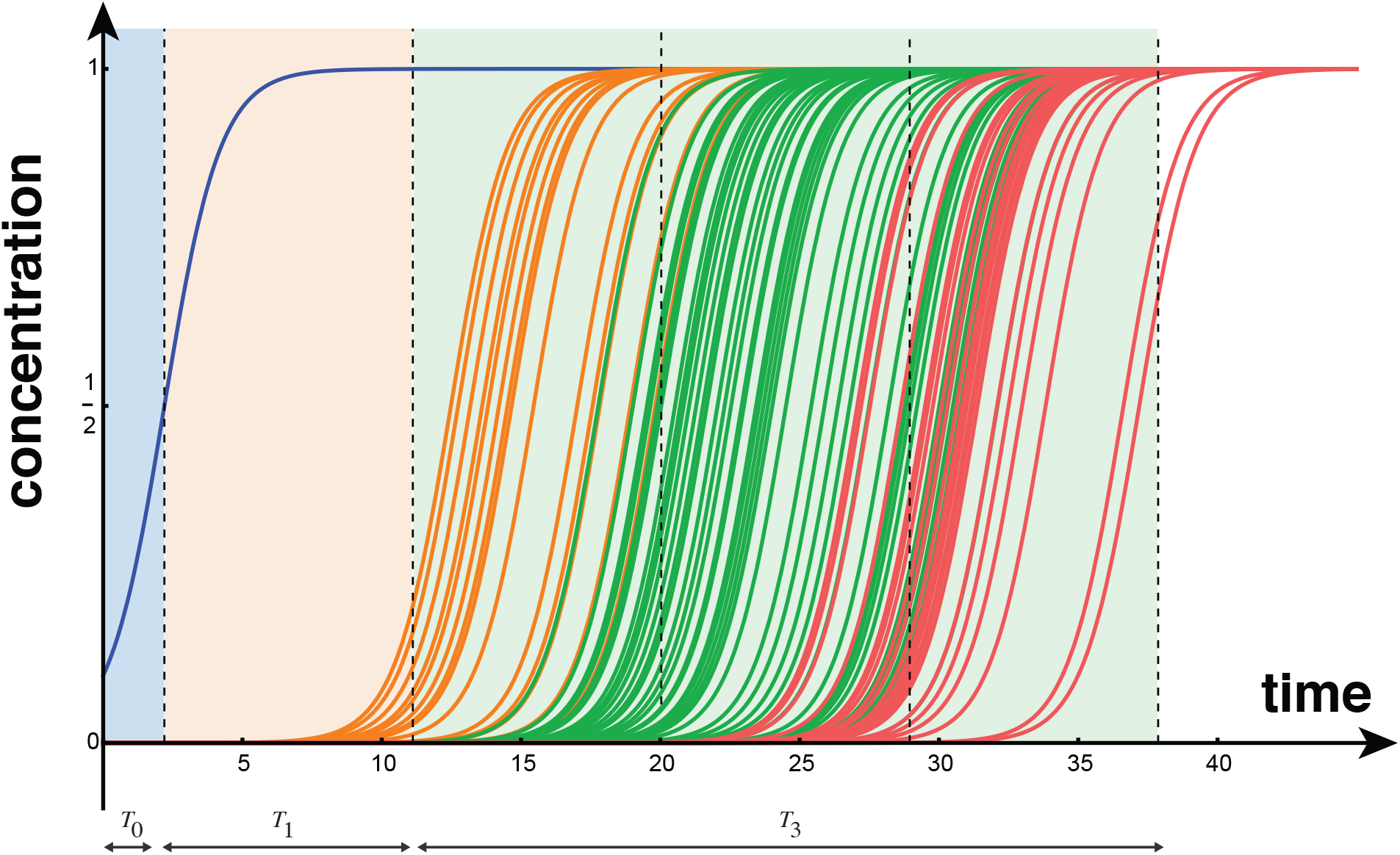
Time scale estimates for the *N* = 83 connectome. Following the local growth at the seeding node with a time scale *T*_0_, the first neighboring is invaded at time *T*_1_. Successive nodes are invaded progressively with a general time scale depending on the number of steps to the last node. Parameter: *α* = 1/year, *ρ* = 0.001/year. Initial condition: *p*_27_ = 1*/*10 and *p*_*i*_ = 0 for all other *i* ∈ {1, …, *N*}. The colors correspond to *γ* as defined in the text: the number of steps from the seed according to the Lambert metric (0: blue, 1: orange, 2: green, 3: red).

The second important time scale is the arrival time at a neighboring node. For a given seed at node *s*, it is given by

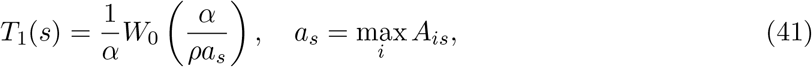

where *a*_*s*_ is the maximum of the *s*th column vector of matrix **A**. If the entries of the weighted adjacency matrix, **A**, are normalized by the maximal entry then, over all possible seeds, the quantity max_*i*_ *A*_*is*_ is, by construction, equal to one. In this case, we can define, for the entire network, the time scale

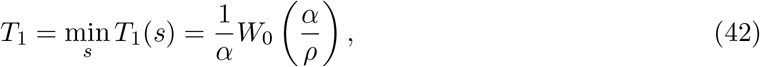

as the *denial period*. The *denial period* (42) corresponds to a time scale quantifying the initial lag in disease invasion from any epicenter. Before the denial period has lapsed, the disease appears to be located at the source and its propagation to neighboring nodes is not yet measurable. The denial period only depends on local connectivity and does not rely on the global, possibly complex, topological characteristics of the network.

If a seeding site is known, or suspected, the local denial period gives a good estimate of the time before an invasion is detectable in nearby regions. The denial period estimates are particularly interesting in the case of modeling Alzheimer’s disease and may provide a window into the early phases of progression for both a typically-assumed entorhinal cortex seeding [38, 34, 39, 40] and the potential alternative seeding sites referenced in connected to non-standard Alzheimer subtypes [41]. Conversely, the global denial period (42) represents an estimate of the minimal window of time necessary for any toxic protein to spread from any seeding location within the brain and is a property of the connectome graph and the choice of weights in the matrix **A**.

The third time scale corresponds to the rapid succession of invasion fronts as shown in Fig. 10. We define, for *i >* 1, *T*_*i*_(*s*) = *iT*_1_(*s*) as typical scale at which nodes that have values *i* with respect to the step distance defined in Section 3, are affected by an initial seeding in node *s*. For instance, the green curves in Fig. 10 characterize nodes whose shortest path to the seeding node has two edges. We see from Fig. 10 that for this network, it is the edge distance that mostly defines the arrival time.

More generally, for a network model over which the system (1) is evolved, the important time scales are *T*_loc_ for the initial growth, *T*_1_ for the first initial spread, and the *pandemic period* where most of the graph has been invaded:

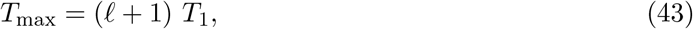

where, *l* is the maximal number of edges between any two nodes in the network defined by **A**. These time scales are independent of the seeding node and the initial condition. These estimates can be further refined for a given initial seed by replacing *T*_1_ by *T*_1_(*s*). For instance, for the *N* = 83 connectome, we have, *l* = 3, setting *α* = 1 year^-1^, *s* = 27, and *T*_max_ = 35 years suggests taking the diffusion constant to be

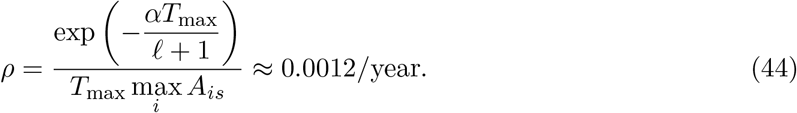

### 5.2 Arrival times and Braak staging

Despite the overall simplicity of the connectome Fisher-KPP model, its dynamics captures well the generalized progression of tau protein neurofibrillary tangles observed in post-mortem studies. In particular, histopathological studies have noted that neurofibrillary tangle follow a six-stage sequence [42, 43], called the *six Braak stages* [44]. While generalized progression sequences have also been proposed [45, 46], here we limit ourselves to the six Braak regions used in the standardized positron emission tomography AV-1451 data pipeline [47] of the Alzheimer’s Disease Neuroimaging Initiative. In line with histopathological observation [42, 43] we expect a progressive ordering of arrival, of tau concentration in different regions, starting in Braak region I corresponding to the entorhinal cortex. Therefore, we solve (1) on a brain connectome with an initial seeding in the bilateral entorhinal cortex. Fig. 8 shows the dynamics of an example simulation, with illustrative parameters in the growth dominated regime, using the lowest resolution *N* = 83 connectome. In this case, the entorhinal seeding sites correspond to nodes 27 and 68.

**Figure 8:**
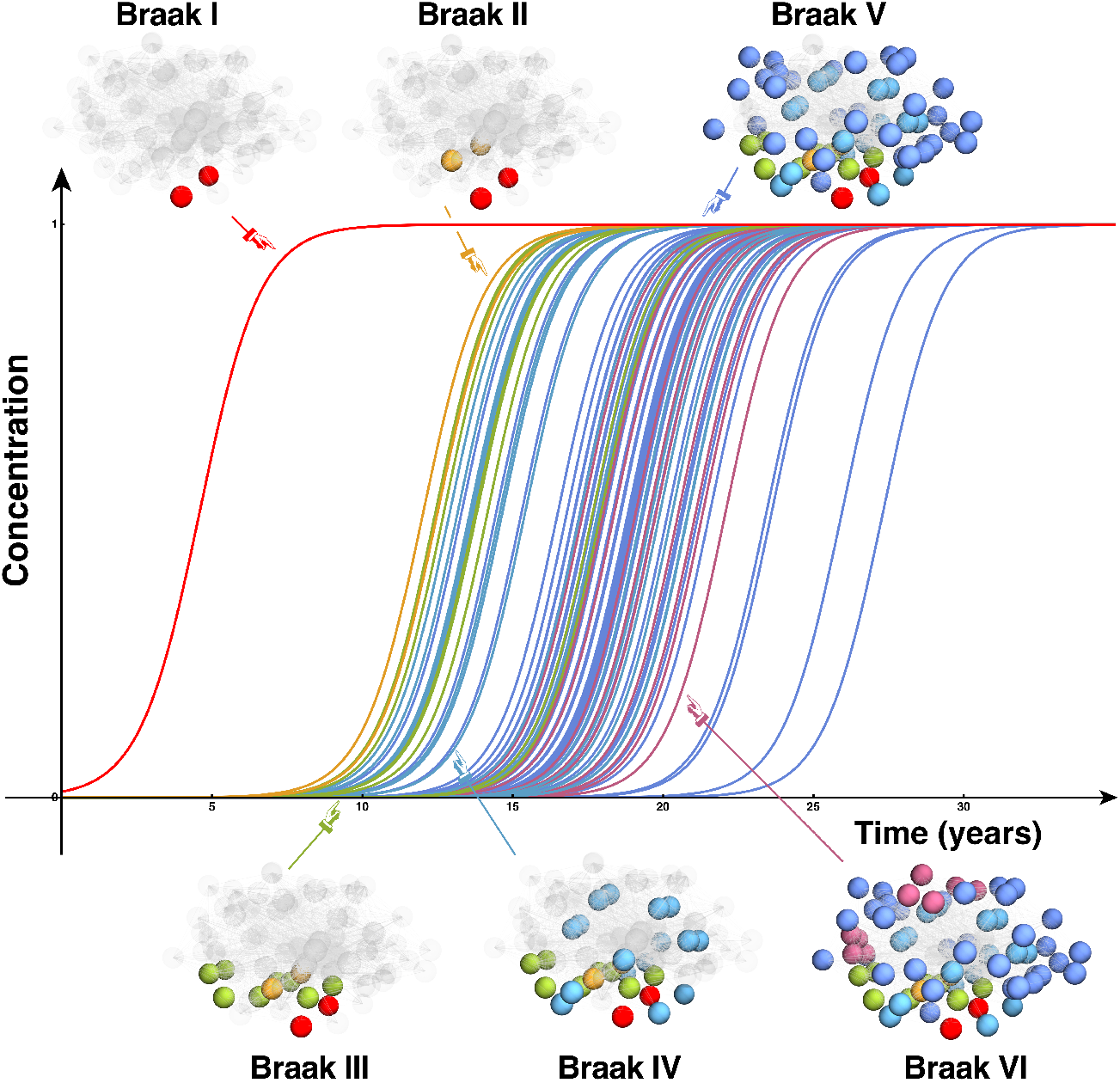
Starting at the entorhinal cortex, the dynamics of the network Fisher-KPP equations recovers some of the key aspects of Braak staging *N* = 83, *α* = 1/year, *ρ* = 5.64 × 10^*−*3^/year. Initial conditions: *p*_27_ = *p*_68_ = 1*/*100 and *p*_*i*_ = 0 for all other *i* ∈{1, …, *N*}. Each of the six colors corresponds to the cortical regions that comprise a Braak stage. For instance, regions that are part of Braak I are in red.

We apply the three methods for arrival times on this problem for the same parameters as in Fig. 8 and compute the approximate linear solutions. In Fig. 9, we give the three arrival times and compare, for each Braak region, the exact numerical solution (solid curves), with the approximate asymptotic solutions (dashed curves). To visualize the results and the quality of the approximations, we average the concentration for each Braak region (as the sum of the concentration over each node in divided by the number of nodes that region). As expected, the nonlinear solution provides a decent approximation of the solution and the best arrival times with comparative scores (12) given by

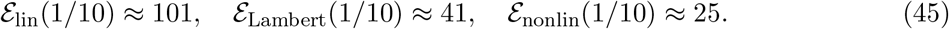

**Figure 9:**
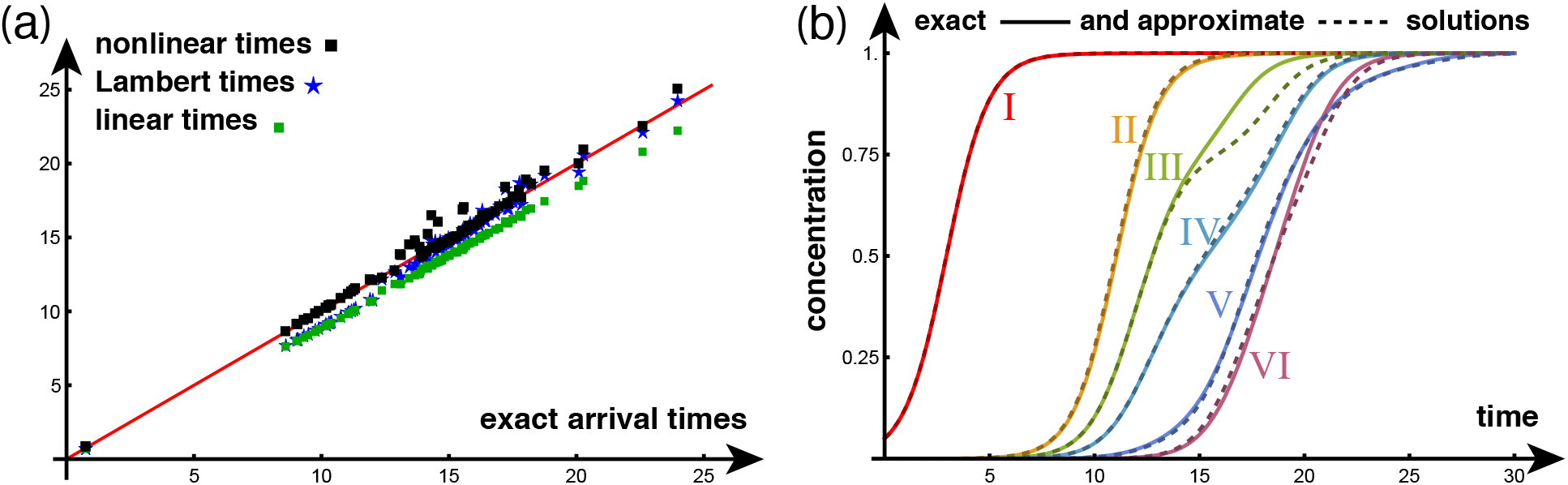
Arrival times at *μ* = 1*/*10 for *N* = 83 connectome (left) and average concentration (right) in each Braak region (parameters and color coding coincide with Fig. 8). Initial conditions chosen so that the total concentration in the entorhinal cortex is 1/10: *p*_27_ = *p*_68_ = 1*/*20 and *p*_*i*_ = 0 for all other *i* ∈ {1, …, *N*}.

The same trends appears, again, in the highest resolution connectome, with 1015 nodes shown in Fig. 10, and the approximate nonlinear solutions provide a good, and best overall, match for both the arrival times and the concentration in the six Braak regions.

**Figure 10:**
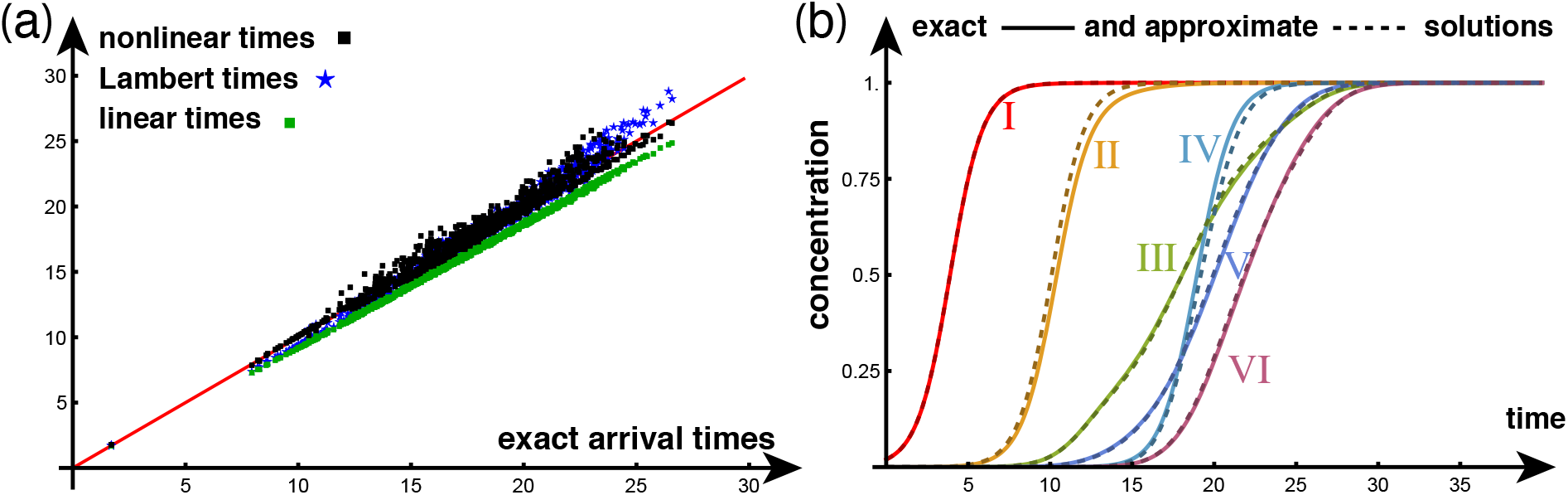
Arrival times at *μ* = 1*/*10 for *N* = 1015 connectome (left) and average concentration (right) in each Braak region (parameters and color coding coincide with Fig. 8). Initial conditions chosen so that the total concentration in the entorhinal cortex is 1/10: *p*_412_ = *p*_413_ = *p*_917_ = *p*_918_ = *p*_919_ = 1*/*50 and *p*_*i*_ = 0 for all other *i* ∈ {1, …, *N*}.

### 5.3 Braid diagrams and braid surfaces

We have applied each approximation method on the lowest and higher resolution connectomes for particular values and are interested in assessing the performance of our methods systematically through staging analysis in the growth-dominated regime (*ρ/α* ≪ 1) as introduced in Putra et al [26]. The idea is to only consider the ordering of the arrival times rather than their numerical values. Following [26], we can plot stagings, across all possible values of the threshold *μ*, by using a *braid diagram*. By way of illustration, Fig. 11(a) shows a simple example of a braid diagram for our exemplar 5-node network given in Fig.1. The braid diagram displays for each value of *μ*, starting at *μ* = *β* = 1*/*100, the order at arrival. For *μ* = 1*/*100, the staging is (3, 4, 2, 1, 5) indicating that node three is the first one at which the concentration reaches 1*/*100 followed by node four, then two, one and five. For higher threshold, the ordering changes. For instance at *μ* = 1*/*2, the ordering is (4, 3, 2, 1, 5). A *braid surface* summarizes a parametric family of braid diagrams. To capture this variability visually, we introduce the notion of *braid surface* that allows for representing multiple braid diagrams associated with different values of *ρ/α* [26]. The idea is to assign a single integer value, in [1, *N* !], and to associate a color label to that integer. For our example, the braid surface in Fig. 11(b) gives eight possible staging depending on both threshold and parameters. Braid surfaces are a powerful means of summarizing staging on a general graph and have been used [26] to show that variations in parameters, connectome resolution, tractography methods and thresholding can cause significant differences in observed Braak staging.

**Figure 11:**
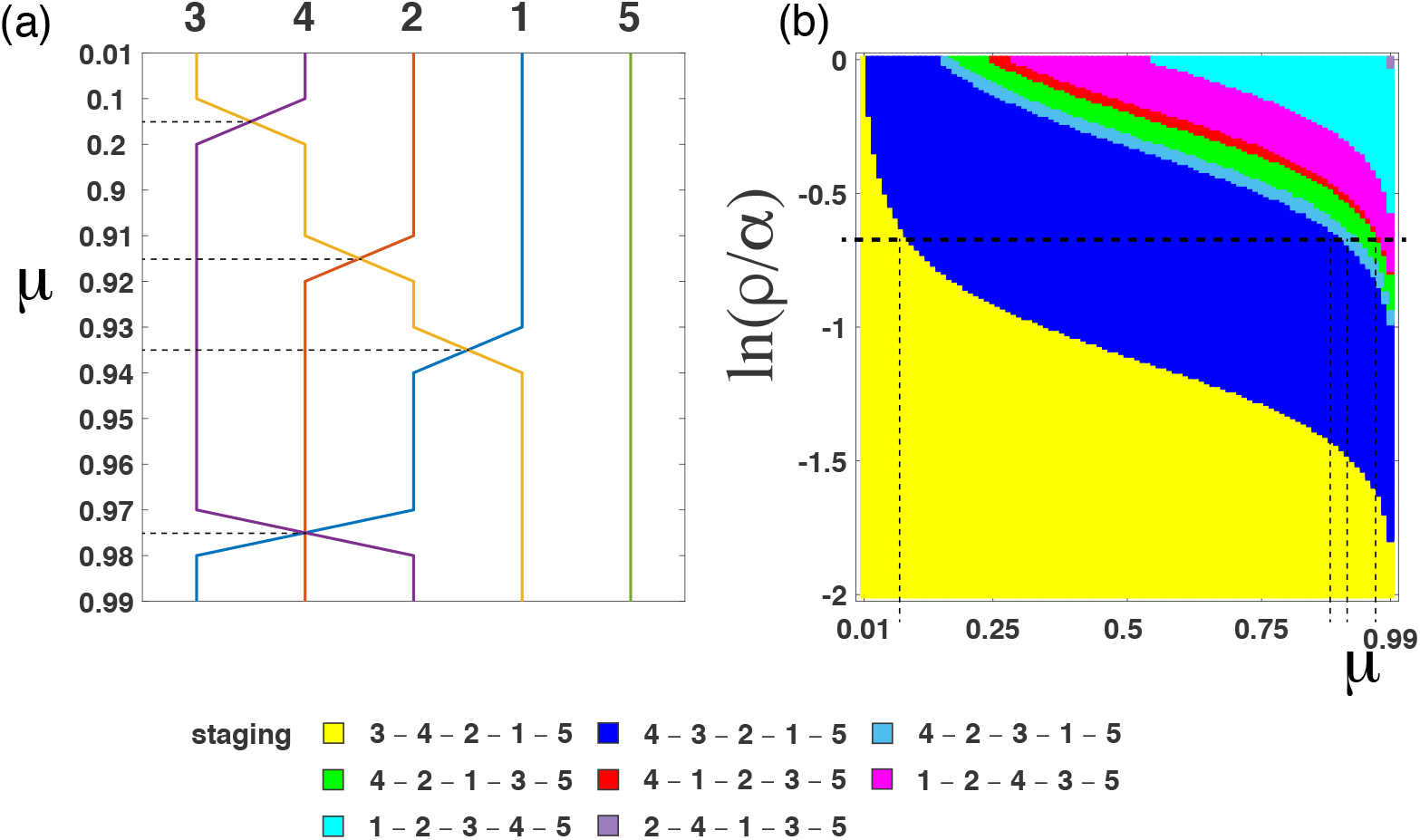
(a) A braid diagram generated using a graph in Fig. 1(a) where ln (*ρ/α*) = −0.7. Dashed horizontal lines highlight examples of a change in staging from one threshold to the next. (b) A braid surface showing staging dynamics as parameter *ρ/α* varies (with a unique seed *p*_3_(0) = *β* = 1*/*100)

Constructing a braid surface can be a computationally expensive task. Suppose that *μ* and log(*ρ/α*) take values in an interval *I*. Determining a braid surface, corresponding to this value range, by exact means requires the discretization of *I* into *M* and *R* subintervals, for *μ* and log(*ρ/α*) respectively, and *M* × *R* solutions of the stiff Fisher-KPP system. The arrival times can also be approximated by our methods discussed. In practice, this trades the computational task of solving a stiff system for the task of solving the problems associated to the estimation methods. Of the three methods, the Lambert method is the most computationally attractive. The Lambert method involves computing the Lambert function for each edge in the graph and determining a shortest path; both tasks have highly performant implementations in many common computing software packages. Moreover, edge distances and shortest paths can be reused for all values of *μ*.

To test the use of the Lambert distance to compute the braid surfaces, we use a the *Kendall tau* distance [48] to measure the distance between two stagings as the minimum number of transpositions necessary to transform one staging into the other. For instance, the Kendall tau distance between staging (1,2,3) and (1,3,2) is *d*_*K*_((1, 2, 3), (1, 3, 2)) = 1, since transposing the last two values of (1, 3, 2) brings the stagings into agreement. Using this notion of distance, we can assess the errors between a braid surface computed explicitly, using (1), and those computed using the linear, Lambert and non-linear estimation methods. Fig. 12(a) shows an eaxct braid surface corresponding to Braak staging on the *N* = 83 node connectome; the yellow color indicates the canonical Braak staging sequence for tau proteins as described by Braak [42, 43]. Figures 12(b)-(d) show the Kendall Tau distance between the exact Braid surface and the Braid surfaces constructed using the three arrival time estimates. As in Fig. 8b, the non-linear approximation (Fig. 12(d)) has the lowest distribution of error, followed by the Lambert distance (Fig. 12(c)) and the linear estimate (Fig. 12(b)). The errors are generally the most pronounced in the diffusion dominated region, where log(*ρ/α*) ≈ 0, and the accuracy of all methods tends to improve with a decrease in the threshold *μ*. The methods for arrival times have been designed under the general assumptions that the threshold is small and the dynamics is growth-dominated. This corresponds to the lower left-hand corned of the braid surface diagram. In that regime, we see that all methods are excellent in recovering the exact staging and can therefore be used systematically for that purpose. These general observations again hold true for the high resolution *N* = 1015 connectome (not shown).

**Figure 12:**
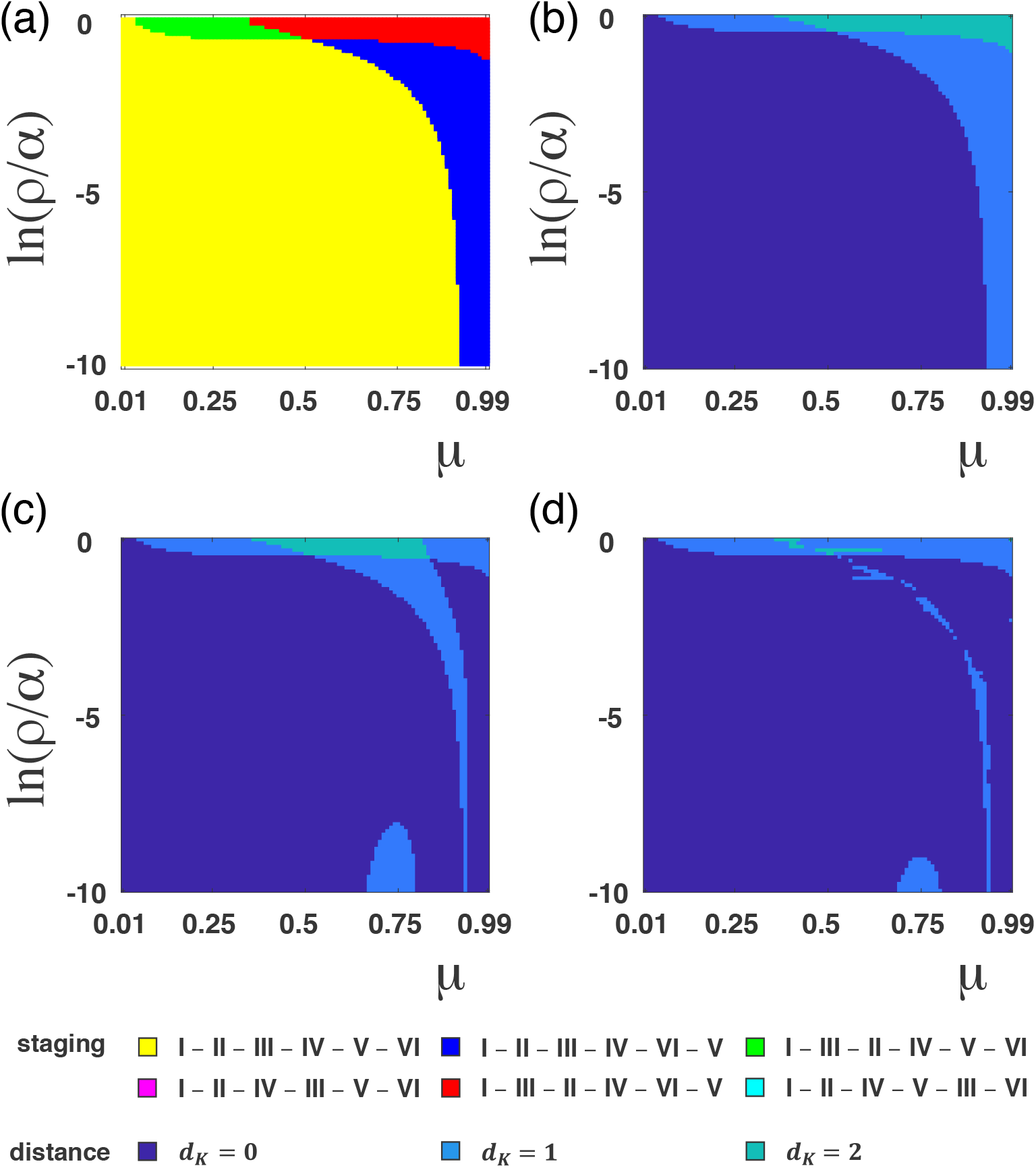
(a) An exact braid surface computed using (1) on the *N* = 83 brain connectome. Kendall Tau errors for the approximated braid surfaces using: (b) the linear method, (c) the Lambert method and (d) the nonlinear method. An initial seeding is chosen so that the average concentration in the entorhinal cortex is less than 1/100: *p*_27_ = *p*_68_ = 1*/*200 and *p*_*i*_ = 0 for all other *i* ∈ 1, …, *N*. Yellow regions correspond to a canonical connectome Braak staging.

## 6 Conclusion

Dynamical processes on a network bring together the combined effects of local dynamics, transport, global topology and interactions between nodes to create complex patterns of evolution. Yet, in the case of a single autocatalytic growth process transported by the graph Laplacian, a simple picture emerges. When diffusion is small enough, compared to the autocatalytic growth, the network is systematically invaded, node by node and the overall process can be characterized by an early seeding and by the arrival time, at each node, of a non-zero concentration.

In this manuscript, we have presented three different and complementary approaches to the problem that provide most of the relevant information about the dynamics. First, we discussed the linear solutions, obtained by a suitable linearization of the entire system of differential equations. We showed that these solutions provide a systematic lower bound for the arrival times. While these estimates have a systematic bias, the method is universal and can be easily generalized to other systems. The linear method also has the advantage of capturing the topology of the entire network through an exponential of the graph Laplacian. The primary drawback of the linear approach is that it requires the numerical solution of a transcendental equation at each node, which may be difficult in the absence of an initial guess.

Second, we introduced the notion of Lambert distance on a graph based on a linear estimate of transport between two neighboring nodes. From the Lambert distance, the arrival times can be estimated by finding the shortest path with respect to this metric. This method is particularly powerful as it is easy to compute and can be used to provide overall approximation of the dynamics. The Lambert method also leads to characteristic time scales, such as the denial and pandemic periods, and endows the network with a metric that is closely associated with the evolving dynamics of interest.

Third, we developed an asymptotic method based on the nonlinear variation of constants. The asymptotic approach builds an approximate solution of the problem that is valid uniformly and has the correct asymptotic properties. This method further improves the Lambert estimates. The primary drawback of this method is that it relies on the existence of an exact solution for the single-node problem. As a result, this approach may fail to generalize to other problems.

Finally, the dynamics and the methods developed herein are motivated by the use of network models in the study of neurodegenerative diseases. Structural connectomes are often characterized by a small-world type topology with highly connected sub-networks and small mean-shortest path length. For the connectomes discussed here, it takes on average less than 3 edges to connect any two nodes in both the *N* = 83 and *N* = 1015 networks. In this context, we have shown that analytical methods to estimate and compute the arrival times are extremely powerful and can capture essential features, such as the correct arrival times and Braak staging.

## Acknowledgment

The work of AG, HO and TT was supported by the Engineering and Physical Sciences Research Council grant EP/R020205/1 to AG. The work of TT was supported partially the John Fell Oxford University Press Research Fund grant 000872 (project code BKD00160) to TT.

## Footnotes

1 The method used here is a bit unorthodox. However, it can be seen as a more or less straightforward generalization of the well-known method of variation of constants for inhomogeneous linear differential equations. As discussed in [27], this method was first proposed by Lagrange in 1788 [28], almost exactly along the lines used here.

